# Synergistic Enzyme Mixtures to Realize Near-Complete Depolymerization in Biodegradable Polymer/Additive Blends

**DOI:** 10.1101/2021.08.25.457667

**Authors:** Christopher DelRe, Boyce Chang, Ivan Jayapurna, Aaron Hall, Ariel Wang, Kyle Zolkin, Ting Xu

## Abstract

Embedding catalysts inside of plastics affords accelerated chemical modification with programmable latency and pathways. Nanoscopically embedded enzymes can lead to near complete degradation of polyesters via chain-end mediated processive depolymerization. The overall degradation rate and pathways have a strong dependence on the morphology of semi-crystalline polyesters. Yet, most studies to date focus on pristine polymers instead of mixtures with additives and other components despite their nearly universal uses in plastic production. Here, additives are introduced to purposely change the morphology of polycaprolactone (PCL) by increasing the bending and twisting of crystalline lamellae. These morphological changes immobilize chain-ends preferentially at the crystalline/amorphous interfaces and limit chain-end accessibility by the embedded processive enzyme. This chain end redistribution reduces the polymer-to-monomer conversion from >95% to less than 50%, causing formation of highly crystalline plastic pieces including microplastics. By synergizing both random chain scission and processive depolymerization, it is feasible to navigate morphological changes in polymer/additive blends and to achieve near complete depolymerization. The random scission enzymes in the amorphous domains create new chain ends that are subsequently bound and depolymerized by processive enzymes. Present studies further highlight the importance to consider host polymer morphological effects on the reactions catalyzed by embedded catalytic species.

## Main Text

Returning plastics back to small molecules generates value-added by-products that can be used for chemical feedstocks and/or microbial metabolization.^[1]^ Besides establishing environmentally friendly chemical processes to transform polymers, the key figure of merit to realize a circular economy is whether near complete polymer-to-small molecule conversion can be achieved. This is particularly important for single-use plastics since microplastic formation is an unavoidable step when discarded plastic products disintegrate.^[2]^ If plastic degradation is incomplete, microplastic particles will be left behind with long-lasting environmental impact.^[1b, 3]^ Current degradation approaches such as thermolysis and solvolysis create secondary pollution.^[4]^ Emerging polymers with dynamic covalent bonds^[5]^ or periodic backbone cleavage points^[6]^ need to be further developed to soften or eliminate requirements of strongly acidic/basic conditions,^[5]^ high temperatures (>100 °C),^[6]^ or expensive and toxic catalysts.^[7]^ Biodegradable polymers, subject to enzymatic degradation, are readily available and promised to be more environmentally friendly. However, their degradation rate, pathway, and extent of conversion strongly depend on enzyme availability. Unless completely degraded, biodegradable plastics have accelerated microplastic formation.^[8]^

Nanoscopically dispersed enzymes inside of semicrystalline polyesters,^[9]^ such as poly(caprolactone) (PCL) and poly(lactic acid) (PLA), can enable near complete (>95%) depolymerization under mild conditions (water at 30-50 °C).^[10]^ The technical key is to bias the enzymatic pathway to degrade polyesters mainly via chain-end mediated processive depolymerization. This bias relies on selective binding of enzymes to the polymer chain ends. After binding, processive enzymes thread polymer substrates through their active site to carry out successive cleavage steps without releasing the chain.^[7, 9a, 11]^ The enzymes’ active site characteristics and the polymer chain arrangements govern the thermodynamic driving force and kinetic pathway of macroscopic plastic degradation. For instance, enzymes that have deep binding domains with fairly large hydrophobic surface areas facilitate processivity by allowing a polymer chain to bind and slide without dissociating.^[9a, 12]^ Local inter-segmental interactions within the polymer’s crystalline lamellae are key factors governing the degradation latency and kinetic rate of biocatalysis.^[13]^ Nevertheless, there is a significant knowledge gap in biocatalysis inside of enzyme-containing plastics, particularly regarding how the hierarchical structures of host plastics affect molecular actions of embedded enzyme nanoclusters.

To date, studies on enzyme-embedded plastics have been focused on neat polymers.^[8, 14]^ However, commercial polymers are almost exclusively formulated in the blend form to optimize properties and cost effectiveness in plastic manufacturing. Even a small fraction of additives can change the parent polymer’s morphology hierarchically.^[15]^ In semi-crystalline polymers, chain ends are, in general, expelled from crystalline lamellae and concentrate in the amorphous domain.^[16]^ This phenomenon has been used to co-localize chain ends and enzyme nanoclusters to facilitate chain-end binding.^[10]^ However, with additive-induced morphological changes, some chain end segments may be immobilized at or near the crystal-amorphous interface to alleviate density anomalies in the adjacent amorphous domain.^[17]^ Interfacial chain ends become more abundant when the crystalline lamellae are bent or twisted, which is promoted by the presence of enzyme protectants (**Figure 1a**, banded spherulites). These chain ends localized at the interfacial area are likely inaccessible for enzyme binding given their restricted mobility and geometric constraints. This may limit processive depolymerization given that enzymatic binding to the polymer chain ends is a rate limiting factor.^[18]^ Thus, it becomes essential to create chain ends in regions accessible to processive enzymes to initiate biocatalysis, as shown in **Figure 1b**. Given the ubiquity of additives in commercial^[19]^ and biodegradable plastics,^[10, 20]^ it is crucial to systematically investigate how additive-induced host morphological changes affect polymer substrate binding, degradation kinetics, and final extent of depolymerization.

**Figure 1.**
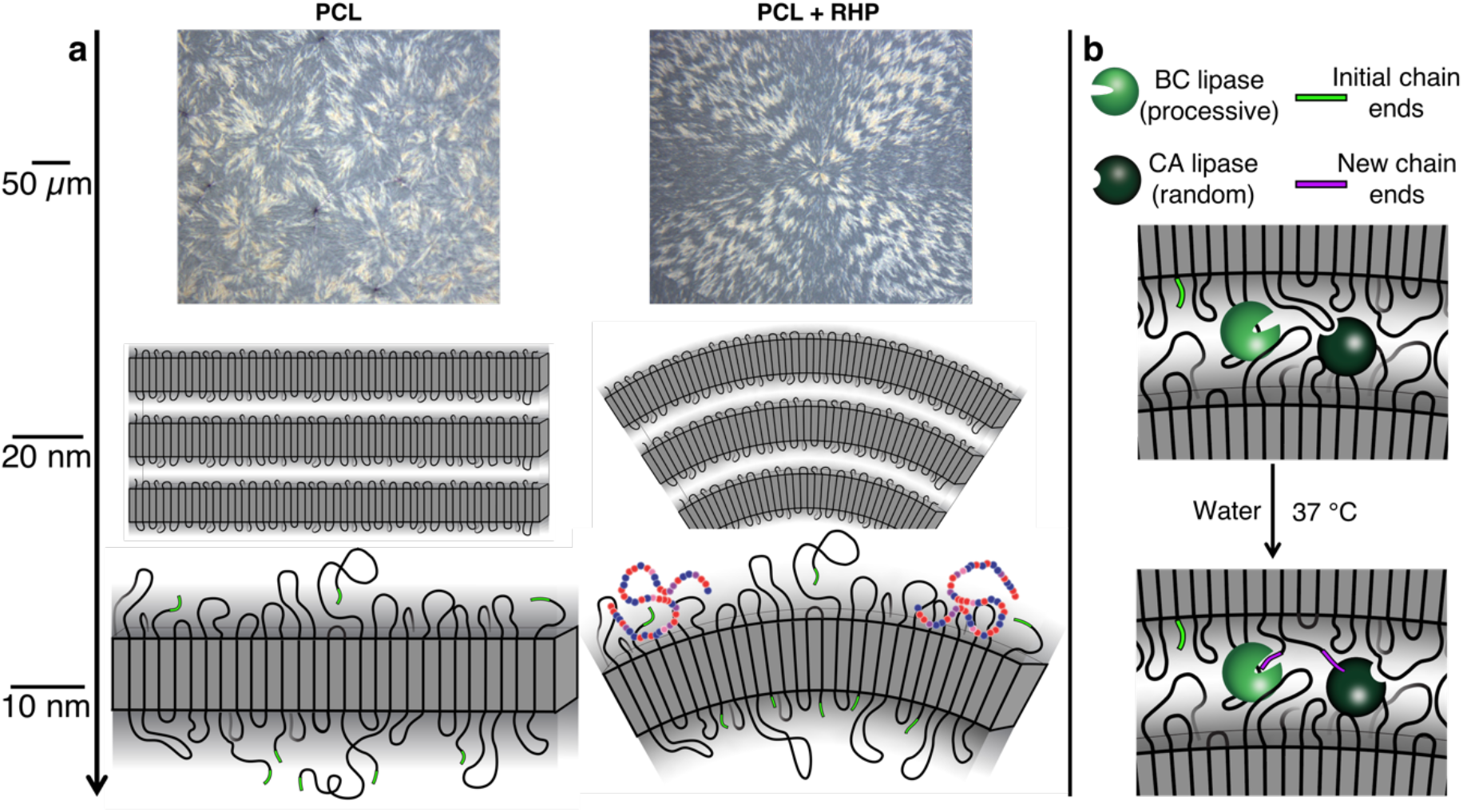
Schematic representation of synergistic enzyme mechanisms overcoming additive-induced inaccessibility of chain ends. (a, top) Polarized optical micrograph of PCL (left) and PCL-RHP_20kDa_ (right) crystallized at 49°C; (a, middle) schematic of lamellae distributions for the respective plastics, as supported by optical and electron microscopy; (a, bottom) schematic highlight the proposed mechanism of lamellae bending caused by transient interactions with RHPs; (b) schematic demonstrating how synergistic mechanisms facilitate complete depolymerization by embedded enzymes—a random scission enzyme can create new chain ends, which can subsequently be bound and depolymerized by a processive enzyme.

Here, using blends of PCL, lipase, and enzyme protectants as examples, we investigated how additives alter the host semicrystalline polyester’s structural arrangement and result in incomplete enzymatic degradation. These molecular insights led to identification of rate limiting factor of biocatalysis and rational design of enzyme mixtures to achieve near complete polymer-to-small molecule conversion. Embedding two enzymes with processive and random chain scission mechanisms, respectively, makes it feasible to create new accessible chain ends to initiate processive depolymerization. These studies demonstrate how additive-induced morphological changes can propagate across hierarchical length scales and offers mechanistic insights into how synergistic enzymes with different mechanisms to modulate the extent of host polymer degradation.

Random heteropolymers (RHPs)^[20b]^ previously designed to stabilize and disperse enzymes are used to embed the enzymes in polymeric matrices. To reduce the blend complexity, two RHPs are used with different molecular weights (MW)—20 kDa (RHP_20k_) and 70 kDa (RHP_70k_)—but the same monomer compositions (**Supporting Information**). Both RHPs are effective to stabilize and disperse hydrolase enzymes.^[20a]^ The PCL:RHP:enzyme mass ratios are also kept the same so that the weight fraction of each component remains constant for all enzyme-containing blends (98.6 wt% PCL, 1.4 wt% RHP, and 0.02 wt% Lipase). Two lipase enzymes are used: *Burkholderia cepacia* (BC-lipase) degrades PCL via processive depolymerization and *Candida Antarctica* Lipase B (CA-lipase) degrades PCL via random chain scission (Figure S1).^[10]^ With lower molecular weight, the number of RHP_20k_ chains is much higher than RHP_70k_ chains. It is expected that RHP_20k_ chains not associated with enzymes are dispersed in the PCL matrix and may affect PCL crystallization due to favorable PCL/RHP interactions.^[15b]^

RHP additives with low MW greatly modify the host matrix’s hierarchical structure. Solution cast PCL and PCL-RHP_70k_ films have primarily straight lamellae bundles; however, lamellae of PCL-RHP_20k_ exhibit substantial bending and twisting, with radii of curvature ranging primarily between 100 and 600 nm (**Figure 2a** and **Figure S2**). The same respective morphologies were observed when RHP-BC-lipase nanoclusters, instead of just RHP, were embedded inside the film (**Figure S3**). PCL was slowly recrystallized from the melt at 49 °C in order to quantify how each RHP may influence the semicrystalline microstructure and growth rate. In polarized optical microscope (POM) images, pure PCL had a simple extinction cross pattern with no defined texture (indicative of straight lamellae bundles) but both RHPs induced a banded extinction cross pattern (**Figure 2b**). Banding in spherulites occurs due to twisting of crystallographic orientation, which arises from twisting and S-shaped bending of lamellae.^[21]^ The band structure for RHP_70k_- containing blend is continuous whereas there is significant irregularity within individual bands for RHP_20k_-containing blend (**Figure 2b insets**). This morphology suggests a higher frequency of twisting/bending in PCL with low MW RHP and is consistent with trends for solution cast films. RHP_20k_ also increases PCL’s crystal growth rate by ∼1.75x during slow crystallization, while RHP_70k_ has negligible effects (**Figure 2c**). These results further demonstrate the ability of RHP_20k_ to modify matrix microstructures.

**Figure 2.**
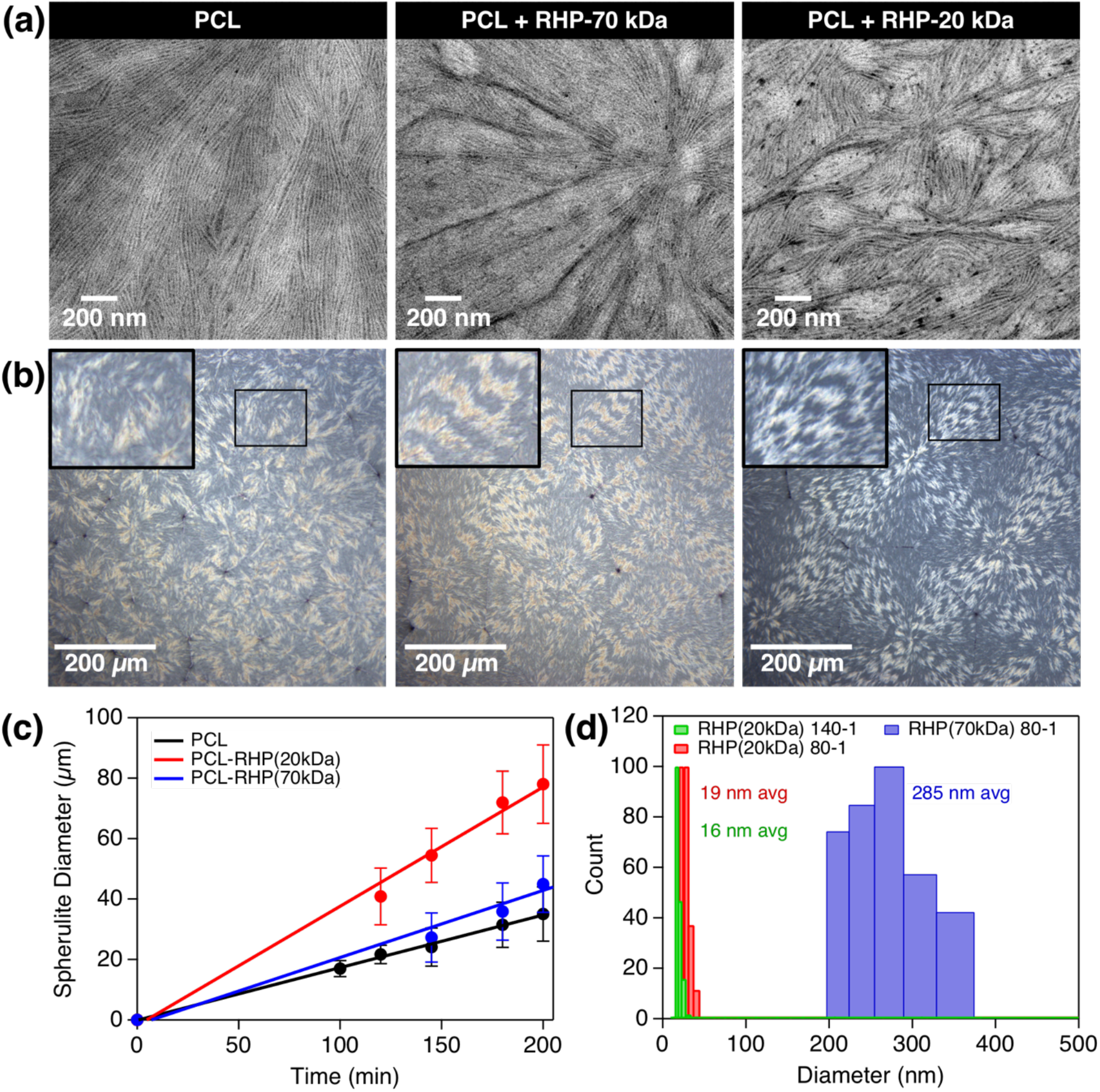
Enzyme protectant (RHP) can alter polyester crystalline morphology. a) TEM images of solution-cast PCL (left), PCL-RHP_70k_-BC-lipase (middle), and PCL-RHP_20k_-BC-lipase (right); B) Polarized optical microscope images of films that were recrystallized from the melt at Tc = 49 °C; c) growth rate of PCL semicrystalline spherulites grown at Tc = 49 °C in the presence of RHP_70k_, RHP_20k_, or no RHP; d) DLS data of BC-lipase protected in toluene by RHP_70k_ or RHP_20k_ at an 80-1 mass ratio; the addition of more RHP_20k_ to achieve a 140-1 mass ratio does not significantly alter the particle size, suggesting that excess RHP molecules likely are responsible for altered PCL matrix properties in solution-cast films.

The mechanism behind the changes in crystalline structures is likely due to transient adsorption of RHPs to nascent crystalline surfaces since RHPs favorably interact with PCL. The RHP additives modify the surface free energy and/or induce imbalanced stresses, causing the lamellae to twist and bend during crystallization.^[15b, 22]^ The reliance of matrix modification on additive MW may be caused by RHP chain mobility in the matrix. RHP_20k_ has fewer entanglements per chain than RHP_70k_, and thus can more easily and rapidly adjust its conformation to interact with nascent crystalline surfaces. RHP distribution within the matrix, which is determined in part by its interactions with BC-lipase, can also influence its interactions with PCL lamellae. The number of possible RHP-enzyme contact points increases with increasing RHP MW. The RHP-BC-lipase clusters are ∼one order of magnitude larger for RHP_70k_ than for RHP_20k_ due to bridging effects (**Figure 2d**). Thus, the amount of free RHP_20k_ within the PCL matrix and the total contact area between RHP_20k_-BC-lipase and PCL should be higher, resulting in more drastic microstructure changes. Thus, inherent properties (i.e., MW) and the distribution of polymer additives in the matrix influence their ability to alter the matrix microstructure.

The semicrystalline microstructure differences associated with different RHP MW have a profound effect on processive depolymerization. Although the depolymerization rates are indistinguishable at early timepoints (≤3 hours) (**Figure S4**), PCL-RHP_20k_-BC-lipase completely stops degrading around 3 hours after ∼65% mass loss while PCL-RHP_70k_-BC-lipase continues to degrade to near-completion (up to 98%) in 24 hours (**Figure 3a**). The PCL-RHP_20k_-BC-lipase film is mechanically fragile but remains intact after depolymerization stops (**Figure 3b**). This ultimately led to formation of microplastics. However, the embedded enzyme remains in the recalcitrant film (i.e., the film that stopped degrading) (**Figure S5a**) and is highly active. Catalytic assays confirmed that the remaining enzymes can hydrolyze a small molecule ester after the film is rinsed and assayed in a separate vial (**Figure S5b**), ruling out enzyme leaching or denaturation to explain recalcitrance from the RHP_20k_ films. We’ve previously shown that crystalline lamellae thickness is a key parameter for processive depolymerization by embedded enzyme. Above a critical lamellae thickness, depolymerization does not occur because the local inter-segmental forces exceed the forces that drive polymer chain threading through the active site. However, DSC confirms that the percent crystallinity and crystalline lamellae thickness are indistinguishable for films made with RHP_20k_ and RHP_70k_ both before and after depolymerization occurs (**Figure 3c**). Furthermore, SAXS confirms that the crystal-amorphous long period is also the same regardless of RHP MW (**Figure 3d**). These experiments characterizing the bulk semicrystalline properties rule out lamellae thickening as a main source of recalcitrance for RHP_20k_ samples. Since the enzyme remains active inside recalcitrant films and the bulk crystalline properties are indistinguishable from films that degrade to near-completion, the recalcitrance arising from RHP_20k_ must be due to subtle molecular-level effects. It is likely that the observable hierarchical microstructure differences may modulate enzyme-polymer substrate interactions.

**Figure 3.**
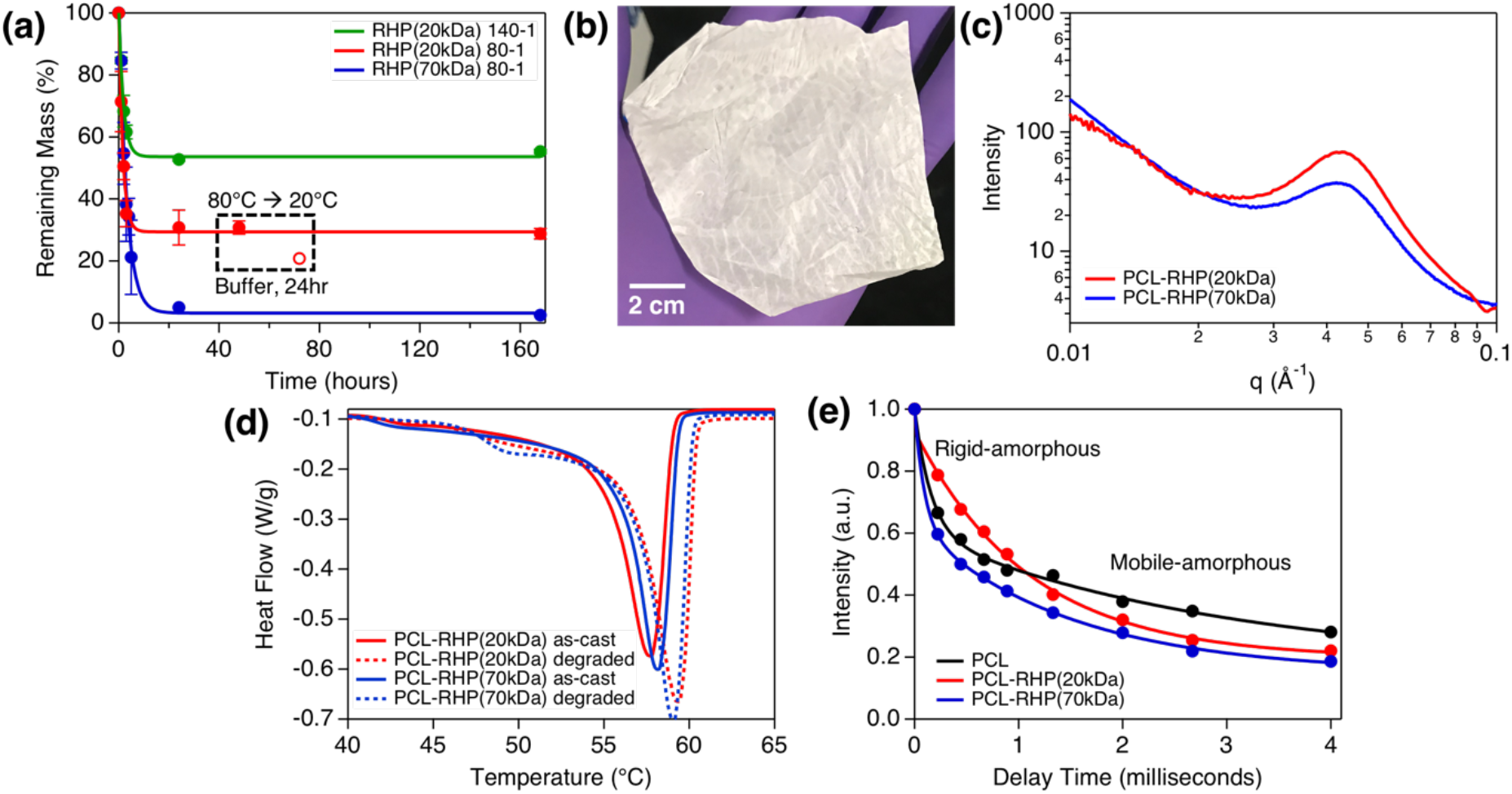
Protectant molecular weight affects extent of depolymerization by embedded processive enzyme. a) Degradation profile of PCL and BC-lipase that was embedded with either RHP(20kDa) or RHP(70kDa); the dotted box shows results from an experiment where films were removed from buffer after they stopped degrading, dried, melted, and recrystallized at room temperature in order to “reset” the crystalline morphology to probe whether chain ends can become accessible again to the enzyme; b) Picture of PCL-RHP(20kDa)-BC-lipase after degradation stops (∼50% mass loss); c) SAXS profiles of PCL and BC-lipase that was embedded with either RHP(20kDa) or RHP(70kDa); d) DSC profiles of PCL and BC-lipase that was embedded with either RHP(20kDa) or RHP(70kDa), before (solid lines) and after (dashed lines) ∼60% mass loss via PCL depolymerization; e) solid state NMR data showing T2 relaxation profiles for as-cast pure PCL, PCL-RHP(70kDa)-BC-lipase after ∼50% mass loss, and PCL-RHP(20kDa)-BC-lipase after ∼50% mass loss

Processive depolymerization by BC-lipase seemingly occurs by selective chain-end binding, so it is feasible that the recalcitrance of PCL-RHP_20k_-BC-lipase arises because bent/twisted lamellae cause inaccessibility of PCL chain end segments in the amorphous domains. Solid-state ^13^C NMR is a particularly powerful technique to decouple segmental-level details within amorphous domains of semicrystalline polymers. Specifically, the T2 relaxation decay of rigid-amorphous segments is 1-2 orders of magnitude faster than that of mobile-amorphous segments due to hindered rigid-amorphous mobility.^[23]^ While ∼50% degraded PCL-RHP_70k_-BC-lipase has similar T2 relaxation as pure PCL in both the rigid-amorphous and mobile-amorphous domains, ∼50% degraded PCL-RHP_20k_-BC-lipase has a substantially slower T2 relaxation in the rigid-amorphous domain after degradation (**Figure 3e** and **Table S1**), indicating higher segmental mobility. This higher rigid-amorphous mobility may arise due to chain end segments being preferentially pinned at the crystal-amorphous interface. Chain end segments can release packing frustration at the curved crystalline/amorphous interfaces. This arrangement would enhance the mobility of the bulk chain segments in the rigid-amorphous domain while immobilizing the chain end segments and making them inaccessible for enzyme binding.

We propose that geometric constraints of bent/twisted lamellae support segregation of chain end segments at or near the crystal-amorphous interface in order to alleviate density anomalies. Specifically, it has been shown that the crystal structure of PCL with bent lamellae remains unchanged compared to that of PCL with planar lamellae;^[15]^ as such, the same number (and mass) of chain segments emerges from the crystalline domain for planar and bent lamellae. However, while planar lamellae have constant available volume throughout the amorphous domains, bent lamellae have continuously decreasing available volume at increasing distances below the crystal-amorphous interface (**Figure S6**). Since the mass remains constant and the available volume decreases, bent lamellae could lead to amorphous domains with density values that become larger than has been measured experimentally. In order to alleviate these potential density anomalies, the chain end segments could preferentially segregate at or near the crystal surface in order to locally reduce the total mass emerging from below the bent crystal-amorphous interface. Recent experimental evidence suggests that as much as 90% of chain end segments can be immobilized at or near the crystal-amorphous interface based on entropic considerations^[17c, 24]^ and/or to alleviate density anomalies. ^[17a, 17b]^ Thus, our experimental and theoretical considerations, combined with recent evidence elsewhere, support a mechanism whereby bent lamellae microstructures inhibit polymer substrate accessibility to processive enzymes by pinning the chain end segments at the crystal-amorphous interface, thereby causing recalcitrance.

Several control experiments were run to support our explanation of recalcitrance arising from chain end inaccessibility. Melting a recalcitrant film that has stopped degrading, followed by quench recrystallization at room temperature (<5 minutes total time), “resets” the crystalline morphology. After resetting the crystalline morphology, recalcitrant PCL-RHP_20k_-BC-lipase films are able to continue depolymerizing by another ∼33% from where depolymerization initially stopped (**Figure 3a, dashed box)**, likely because the new crystalline domains formed during recrystallization made some chain end segments available to the remaining enzyme that was previously incapable of accessing them. This experiment is analogous to removing a roadblock for chain end binding, and a similar example has been demonstrated for surface erosion of cellulose via processive cellulases.^[25]^ Furthermore, increasing the concentration of RHP_20k_ in PCL films by 1.75x while keeping enzyme loading constant leads to more substantial recalcitrance, as depolymerization stops after ∼35% mass loss (**Figure 3a, green**). DLS data shows there is no significant change in RHP-BC-lipase cluster size as RHP loading increases by 1.75x (**Figure 2d, green**), supporting that the recalcitrance associated with RHP_20k_ stems from matrix effects rather than differences in RHP-enzyme interactions. Finally, we reduced the PCL MW to increase the percentage of chain ends and then analyzed the depolymerization rate when RHP_70k_-BC-lipase was embedded—higher MW RHP was used to avoid additive-induced recalcitrance. When all PCL MW samples have indistinguishable lamellae thicknesses—controlled by melt recrystallization to remove thermodynamic effects from different local inter-segmental interaction strengths in crystalline lamellae—the lowest MW sample with the highest percentage of chain ends depolymerizes the slowest while the two higher MW samples have similarly faster depolymerization rates (**Figure S7**). Although this experiment can only be semi-quantitative because of challenges with obtaining perfectly identical semicrystalline morphologies across different PCL MW matrices, the general trend has been reproduced and suggests that PCL chain end binding is a rate-limiting step for processive depolymerization. Taken together, our controls support the proposed explanation that RHP_20k_ inhibits PCL chain end accessibility to BC-lipase by causing the lamellae to bend/twist, which drives chain end segments to pin themselves at the crystal-amorphous interface.

We hypothesized that creating new chain ends in the immediate vicinity of the processive enzyme could overcome the additive-induced recalcitrance. Therefore, we embedded a random scission lipase from *Candida antarctica* (CA-lipase) along with BC-lipase and RHP_20k_. When BC-lipase/CA-lipase mixtures are embedded, the depolymerization rate was indistinguishable from that of just BC-lipase from 0-3 hours (**Figure 4a, inset**). However, the CA-plus BC-lipase films continued to slowly degrade over time to near-completion (consistently >90% depolymerization) (**Figure 4a**). This is in direct contrast to the complete recalcitrance after ∼65% mass loss when only BC-lipase was embedded. The depolymerization rate after 3 hours depended on CA-lipase loading, suggesting that the creation and binding of new chain ends is the rate-limiting factor for this synergistic enzyme approach. When the RHP_20k_ concentration is increased by 1.75x with the same CA- and BC-lipase loadings, the rate again slows significantly after ∼3 hours, but depolymerization continues slowly to over 90% (**Figure 4b**). The control experiments showed that depolymerization stops at 35% for just BC-lipase with 1.75x RHP_20k_. It should be noted that when just CA-lipase is embedded in PCL without BC-lipase, depolymerization stops after ∼12% mass loss. This is contributed to the inability of CA-lipase to depolymerize PCL segments in the crystalline domain. Thus, it is reasonable to conclude that the continued depolymerization for the combined enzyme loading is not due to action solely from addition of CA-lipase. Rather, it is from the synergistic effects of creating new binding sites for BC-lipase to processively depolymerize. Furthermore, Gel permeation chromatography (GPC) studies were carried out. Molecular weight analysis of films at several points along the depolymerization pathway shows the synergistic mechanisms in action (**Figure 4c**). Below 3 hours, GPC shows no noticeable shift in MW of the remaining film, consistent with a processive-dominated depolymerization. However, after the degradation rate slows significantly, the MW continually broadens and shifts to longer elution times, characteristic of random chain scission. Together, all results are consistent and support that CA-lipase is in fact creating new chain ends in the remaining film, which BC-lipase then presumably binds to initiate processive depolymerization.

**Figure 4.**
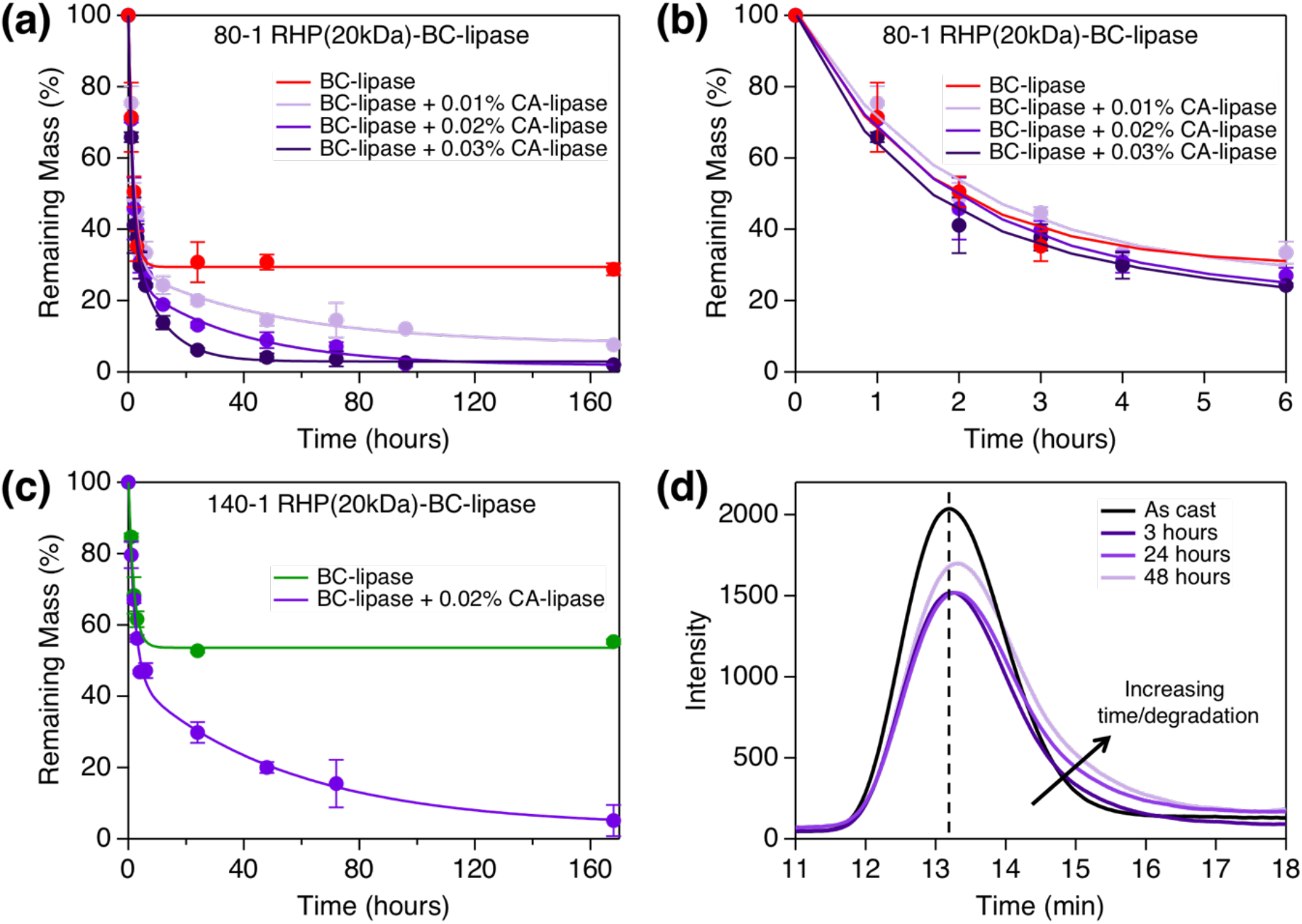
Synergistic enzyme mechanisms overcome protectant-induced recalcitrance. a) Degradation profile of PCL containing BC-lipase with 1.4 wt% RHP(20kDa) (80-1 RHP-BC-lipase ratio) and the specified concentration of CA-lipase; b) Zoomed-in portion of graph from Figure 4a demonstrating that samples with and without CA-lipase have the same degradation rate between 0 and 4 hours; c) Degradation profile of PCL containing just BC-lipase or BC-lipase and CA-lipase embedded with 2.45 wt% of RHP20k (140-1 RHP-BC-lipase ratio); d) GPC profiles of just the remaining PCL film containing BC-lipase and CA-lipase (0.02%) over time.

By using a model biodegradable polyester with embedded enzymes and additives, present studies demonstrate how morphological changes of a semicrystalline polymer’s microstructure can propagate across multiple length scales to alter segmental arrangements and extent of degradation. Specifically, results shown here demonstrated how the additive-driven redistribution of polymer chain ends causes depolymerization recalcitrance when only one type of enzyme was embedded. By synergizing different biocatalysis mechanisms, it became experimentally feasible to modulate the reaction pathways to overcome the observed recalcitrance. This ultimately led to near-complete depolymerization, which is essential for achieving a circular plastics economy. Given the ubiquity of additives in commercial plastics and diverse enzymes available to catalyze similar reactions, there are numerous opportunities to program plastic lifecycle via rational design of enzyme-embedded plastics. Once again, present studies remind us of the importance to investigate catalyzed reactions inside of polymeric matrices and the necessities to evaluate the effect of complex phase behavior of polymer blends and hierarchical nature of semicrystalline polymers. It is our belief that such studies should be carried out for bio- and synthetic catalysts as well as for both degradation and plastic upcycling.

## Experimental Section/Methods

### Materials

The random heteropolymers (RHPs) were synthesized as previously reported.^[20b]^ Both the 70 kDa and 20 kDa RHPs had a molar composition of 50:20:25:5 methyl methacrylate:ethylhexyl methacrylate:oligoethylene glycol methacrylate:sulfopropyl methacrylate (MMA:EHMA:OEGMA:SPMA). PCL (all molecular weights) was purchased from Sigma Aldrich and used without further purification. Amano PS Lipase from *Burkholderia cepacia* was purchased from Sigma Aldrich and purified using previously reported methods.^[26]^ Candida Antarctica Lipase B was also purchased from Sigma Aldrich and used as purchased. Enzyme concentrations in the stock solutions that were eventually used to make RHP-enzyme clusters were confirmed by monitoring the absorbance peak at 280 nm in ultraviolet-visible (UV-vis) spectroscopy.

### Methods

#### Solution casting of PCL-RHP-BC-lipase films

RHP and enzymes were mixed in aqueous solution, flash-frozen in liquid nitrogen and lyophilized overnight to form the RHP-enzyme complexes. The dried mixture was resuspended directly in the PCL solution. RHP was mixed with purified BC-lipase in a mass ratio of 80:1 (total polymer matrix mass =98.6 *wt%*) or 140:1 mass ratio (total polymer matrix mass = 97.6 *wt%*); the BC-lipase concentration in the films, unless otherwise stated, was 0.02 *wt%*.

Unless otherwise specified, PCL with an average MW of 80 kDa was used as the polyester matrix. PCL was dissolved in toluene at 4 wt% concentration and stirred for at least 4 hours at 55°C to ensure complete dissolution. The dried RHP-BC-lipase complexes were resuspended directly in the polymer solution at 0.02 wt% enzyme concentration. Mixtures were vortexed for ∼5 minutes before directly casted on a glass slide and air dried in a chemical fume hood at room temperature.

#### Characterization of PCL-RHP-BC-lipase films

Dynamic light scattering (DLS) was used to obtain the complex’s particle size in toluene. DLS was run on a Brookhaven BI-200SM Light Scattering System using a 90° angle.

Samples were vacuum dried after degradation for 16 hours prior to running SAXS at beamline 7.3.3 at the Advanced Light Source (ALS). X-rays with 1.24 Å wavelength and 2 s exposure times were used. The scattered X-ray intensity distribution was detected using a high-speed Pilatus 2M detector. Images were plotted as intensity (*I*) *vs. q*, where *q* = (4πλ^-1^) sin(θ), λ is the wavelength of the incident X-ray beam, and 2θ is the scattering angle. The sector-average profiles of SAXS patterns were extracted using Igor Pro with the Nika package.

TEM images of solution cast films were taken on a JEOL 1200 microscope at 120 kV accelerating voltage. Vapor from a 0.5 *wt*% ruthenium tetroxide solution was used to stain the RHP-lipase and the amorphous PCL domains. TEM images of curved lamellae were analyzed using ImageJ software. Using the line tool, the ends of the curved lamellae were extended radially until they intersect. The sector was then measured for the central angle and radius of curvature using the angle and measurement tools, respectively. The arclength of each individual lamellae was determined using these measurements.

Polarized optical microscopy (POM) images were taken on a benchtop optical microscope with a polarizer attachment to observe the spherulite morphology. Samples analyzed by POM were recrystallized at 49°C from the melt using a standard temperature-controlled hot stage to enlarge spherulite size because the solution cast films possessed spherulites too small to characterize. Spherulite growth rates of melt processed films were obtained by capturing POM image at specific time intervals and fitting a line to the data.

#### Degradation by BC-lipase

Solution cast films of ∼5 mg were degraded in 2 mL of sodium phosphate buffer (25 mM, pH = 7.2) held in a 37°C water bath. Films were removed from buffer at designated time points, rinsed, dried, and weighed to determine mass loss.

#### Control experiment—small molecule ester assay to confirm embedded BC-lipase is still active after degradation stops

For recalcitrant films, a small molecule ester assay was performed after degradation plateaued to check the activities of complexed lipase enzymes. 4-nitrophenyl butyrate was dissolved in buffer in a separate vial from the one in which depolymerization occurred to rule out effects from enzymes that had entered the buffer from portions of the film that were already depolymerized. These colorimetric assays (substrate is clear, product is yellow) run with 0.5 mM ester solution show that the enzyme remaining in the recalcitrant film remains highly active towards small molecule esters, ruling out enzyme deactivation or leaching as the primary source of recalcitrance and suggested that the remaining enzyme simply does not have access to the polymer matrix substrate.

#### Control experiment—”reset” crystal morphology of recalcitrant films

As a control experiment to confirm that recalcitrance is due to a property of the matrix, films containing RHP_20k_ that stopped degrading were dried, melted at 80°C for 5 minutes, and rapidly recrystallized (∼1 minute) at 20 °C. These films were placed back into phosphate buffer and allowed to continue to degrade under the same set of conditions described above. Depolymerization for the previously recalcitrant films continued after “resetting” the crystalline morphology, supporting the explanation that recalcitrance stems from polymer substrate inaccessibility due to some morphological detail.

#### Differential scanning calorimetry (DSC) to quantify lamellae thickness and percent crystallinity before and after degradation for PCL-BC-lipase with different RHP MWs

Crystallinity of the films before and after degradation were probed using differential scanning calorimetry (DSC). Each PCL sample (∼3-5 mg) was pressed into an aluminum pan, with an empty aluminum pan used as the reference. The temperature was ramped from 25°C to 70°C at a rate of 2 °C min^-1^. The sample’s enthalpy of melting, quantified by integrating under the melting peak, was normalized by the enthalpy of melting for 100% crystalline PCL, 151.7 J g^-1^, to determine percent crystallinity. The crystal-amorphous long period obtained from SAXS combined with the percent crystallinity from DSC, as well as the melting temperature from DSC (which serves as a proxy for crystalline lamellae thickness based on the Thompson-Gibbs relationship^[27]^), both confirm that PCL-RHP-BC-lipase samples have indistinguishable lamellae thickness, long periods, and percent crystallinity.

#### Control experiment—PCL MW with controlled lamellae thickness to probe chain end effects

Melt processing was exploited to obtain PCL films with similar crystalline lamellae thicknesses across three different PCL molecular weights. The films were dropcast from a toluene solution and then melted at 80°C for 5 minutes. The 80 kDa and 45 kDa films were then recrystallized for 1 day at 32°C, and then 10kDa films were recrystallized for 1 day at 30°C. These recrystallization conditions produced PCL-RHP-BC-lipase films with similar crystalline lamellae thicknesses, as determined by monitoring the melting temperature using DSC (Supplementary Figure S6a). Note that films here denoted as “PCL-10 kDa” also have 15% (by mole) of 45 kDa chains added in — the pure 10 kDa film did not have adequate mechanical properties to remain as a freestanding film, so higher MW chains were added to eliminate leaching as a potential concern. “PCL-80 kDa” and “PCL-45 kDa” are comprised solely of the specified molecular weights. Degradation was then carried out as specified above. This experiment was run three separate times using slightly different degradation temperatures (anywhere from 40°C-43°C), and each time the same trends were observed: 45 kDa and 80 kDa samples degraded with similar rates, while 10 kDa samples degraded measurably slower. The data reported here is from degradation at 41°C buffer.

#### Solid-state NMR (to check for mobility in rigid- and mobile-amorphous domains)

##### Data Collection

Solid-state NMR measurements were conducted using a Bruker AV-500 (500 MHz) spectrometer. PCL films were cut into small pieces and subsequently packed into a half rotor. ^13^C chemical shifts were referenced to tetramethylsilane (TMS) using the methylene peak of adamantane (38.48 ppm). All measurements were conducted at room temperature and magic angle spinning (MAS) rate of 9kHz. Recycle delay was set to 2s, which is sufficient for saturation recovery of the amorphous chains (T_1_ = 0.19s) but not the crystalline domains (T_1_ = 28s and 276s). These experimental settings enable us to probe only segmental dynamics in the amorphous domain, which possesses two different T_2_ decay times—the rigid-amorphous segments decay quickly due to constrained mobility, while the mobile-amorphous segments decay slowly (10-100x slower) due to high mobility.^[23]^ NMR ^13^C spin-spin relaxation (T_2_) was obtained using Hahn echo through direct excitation. The experiment included 8 delays synchronized to the MAS rotor period.

##### Data Analysis

T_2_ calculations of PCL were centered on the methylene peak at 64.26 ppm to avoid peak overlap with other groups. Integrals were obtained by line fitting a Gaussian peak from 60-70 ppm. T_2_ relaxation times were obtained by fitting magnetization decay vs delay time to a two-component exponential curve.

#### Calculations to justify hypothetical explanation for chain end segregation to rigid-amorphous domain

To justify the observed chain end redistribution, a basic lattice model was used to calculate the changes in lattice sites for a curved structure. The number of lattice sites per area was taken to be 0.0536 sites angstrom^-2^, based off of previous unit cell characterization.^[23]^ To determine the total number of lattice sites, the constant was multiplied by the surface area of that layer, which differed depending on the geometry. Planar lamellae have constant surface areas while curved structures, modeled as a sphere, have areas that change as a function of the squared radius. For a given curved lattice, the percent of available lattice sites *x* distance below the surface can be approximated by (*r – x*)^2^ *r* ^-2^, where *r* is the approximate radius of the sphere. Supplementary Figure S5 shows the results of varying the radius while maintaining a distance of 4 nm below the crystal/amorphous interface.

#### Degradation mechanisms with synergistic lipases

RHP/BC-lipase/CA-lipase complexes were formed to colocalize BC-lipase and CA-lipase to the same microenvironment within the PCL films. For BC-lipase/CA-lipase co-complexes, the RHP to BC-lipase ratio was kept at 80:1 and different CA-lipase amounts were added to give ratios of 80:0.5, 80:1, or 80:2 RHP:CA-lipase (as specified on the graph in Figure 4a, where the “x” amount is CA-lipase concentration relative to BC-lipase concentration). Samples were lyophilized, dissolved in PCL, and casted into films as previously described. Solution casted films were degraded were degraded in 2 mL sodium phosphate buffer (25 mM, pH = 7.2) held in a 37°C water bath. Films were removed at designated time points, rinsed, dried, and weighed to determine mass loss. For films with higher RHP concentration (Figure 4b), RHP:BC-lipase ratio and RHP:CA-lipase ratio were both 140:1. PCL 80 kDa was used as the matrix for all synergy experiments.

To monitor the PCL MW distribution throughout the degradation pathway by the synergistic enzymes, the remaining films and degradation by-products were lyophilized overnight and resuspended in GPC solvent (THF) following a total concentration of 2 mg mL^-1^ remaining film. 20 uL of solution was injected into an Agilent PolyPore 7.5×300 mm column.

## Supporting Information

Supporting Information is available from the Wiley Online Library or from the author.

## Acknowledgements

This work was supported by the U.S. Department of Energy, Office of Science, Office of Basic Energy Sciences, Materials Sciences and Engineering Division (DOE-BES-MSE) under Contract DE-AC02-05-CH11231, through the Organic−Inorganic Nanocomposites KC3104 program (C.D, B.C Blend morphology, degradation process and NMR studies). The US Department of Defense, Army Research Office support is under contract W911NF2110128 (Enzyme stabilization and biocatalysis). C.D. was partially supported by the National Defense Science and Engineering Graduate (NDSEG) Fellowship. Scattering studies at the Advanced Light Source were supported by the U.S. Department of Energy, Office of Science, Office of Basic Energy Science under contract DE-AC02-05CH11231.

## ToC Text

When enzymes with deep, narrow active sites are embedded inside of plastics, chain end-mediated processive depolymerization is realized. However, polymer additives can induce bending in crystalline lamellae, which causes chain ends to preferentially segregate at the crystal/amorphous interface where they are inaccessible for enzyme binding. Synergistic enzyme mechanisms can be exploited to create new chain ends and facilitate near-complete depolymerization.

## ToC Figure

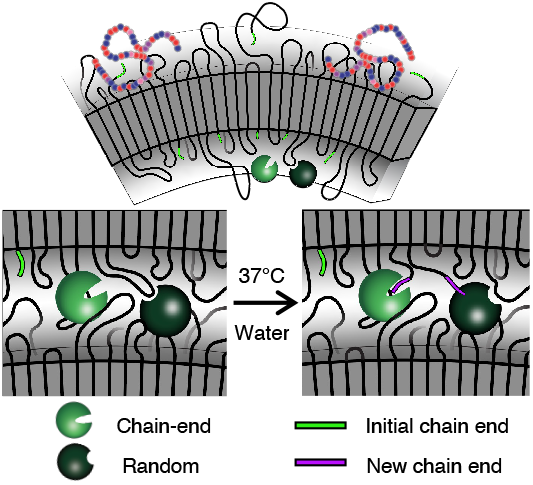

## Supporting Information

**Figure S1.**
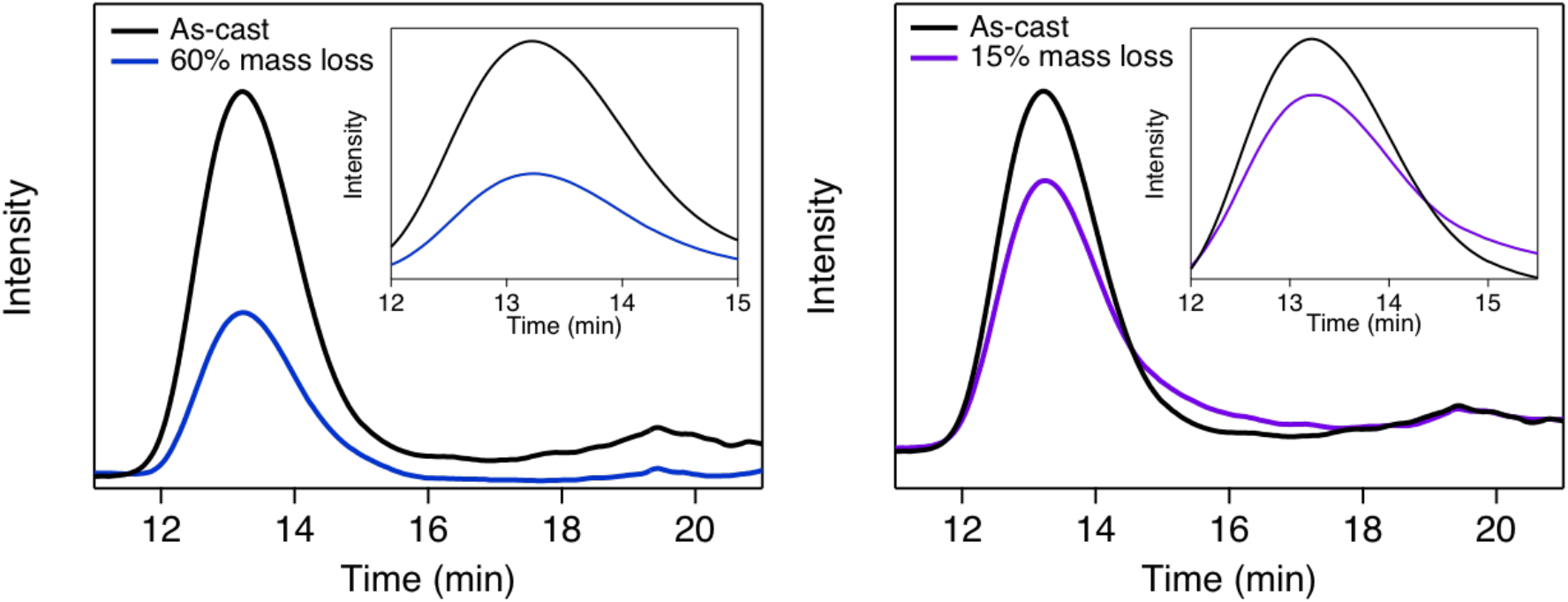
GPC curves of PCL degraded in 37 °C buffer by BC-lipase alone (left) and CA-lipase alone (right) after 60% and 15% mass loss, respectively. RHP_70k_ was used to embed the enzymes in this experiment. Note that PCL containing just CA-lipase only degrades by ∼15% before degradation stops, whereas PCL containing just BC-lipase degrades to near-completion. Insets are zoomed-in portions of relevant portions for each curve.

**Figure S2.**
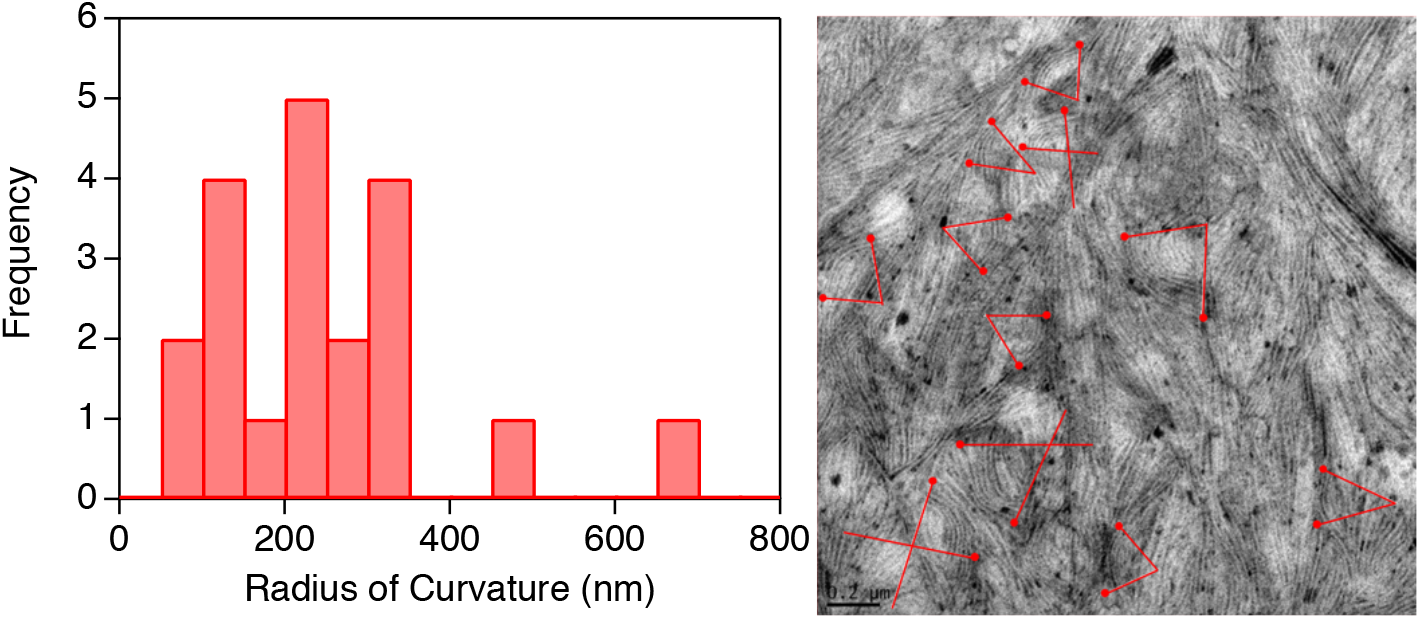
Histogram of measured radii of curvature for PCL-RHP20k from TEM images (left) and example of how curvatures were measured using ImageJ on TEM images (right).

**Figure S3.**
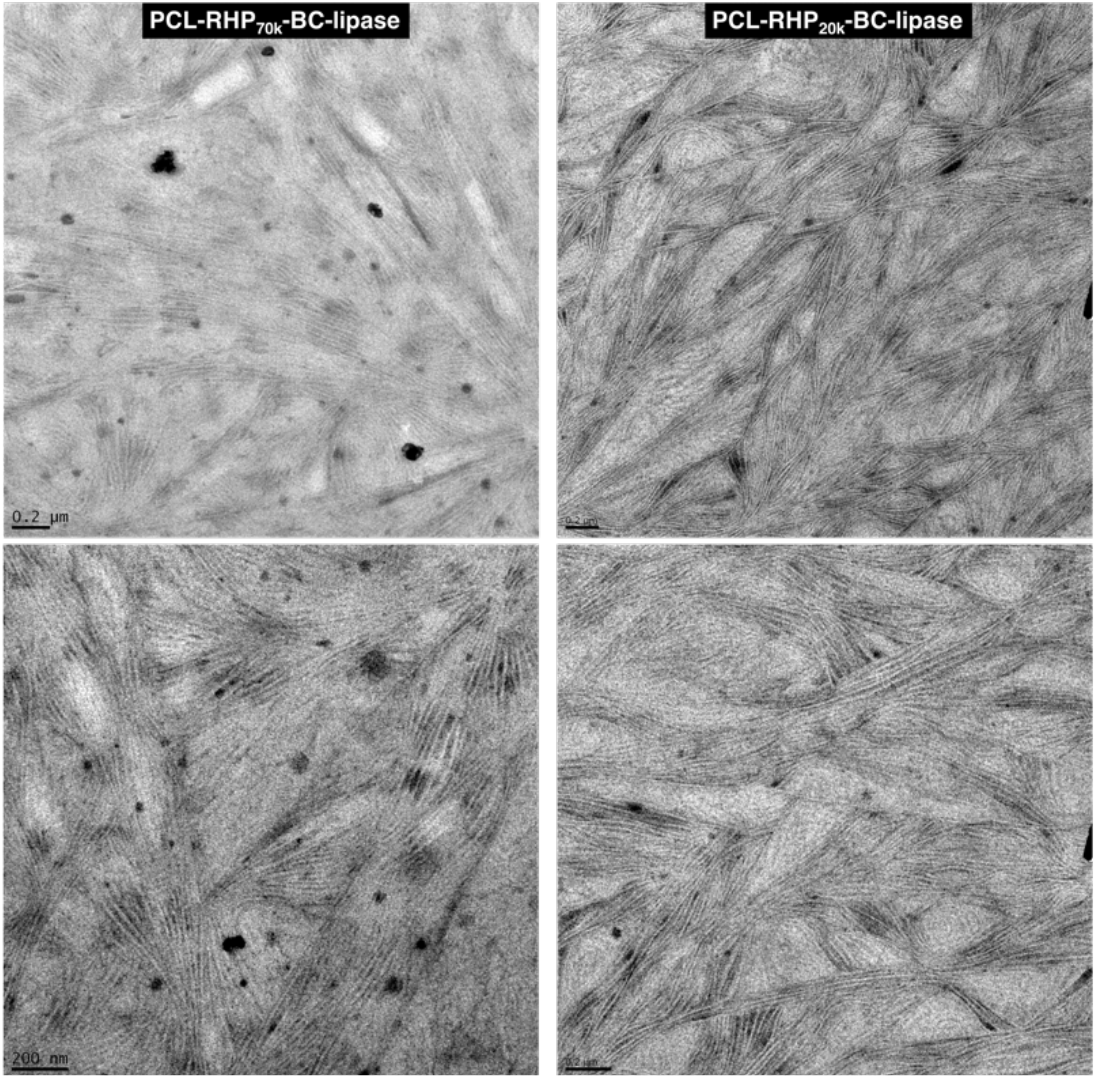
TEM images of PCL-RHP70k-BC-lipase (left) and PCL-RHP20k-BC-lipase (right).

**Figure S4.**
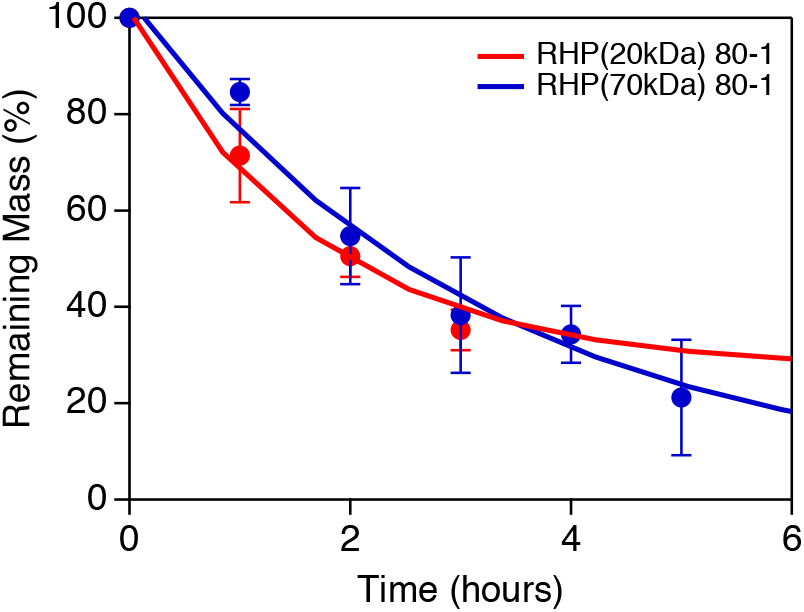
Depolymerization rate from t=0 hours to t=6 hours (enlarged from Figure 3a) of PCL-RHP20k-BC-lipase and PCL-RHP70k-BC-lipase.

**Figure S5.**
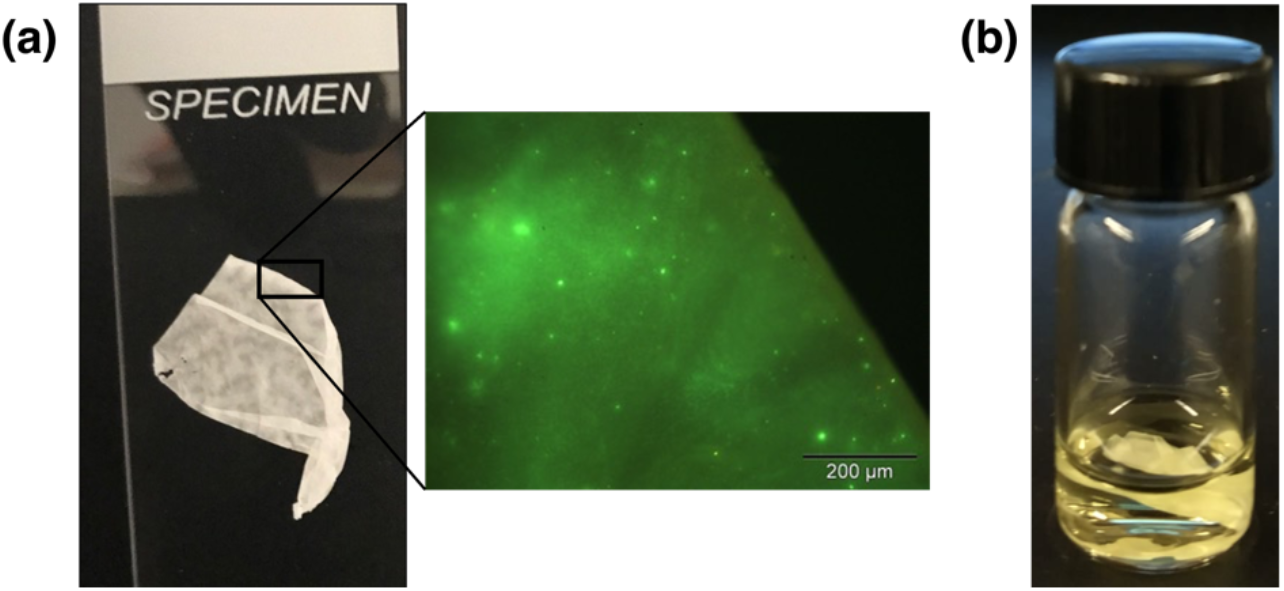
a) Recalcitrant PCL-RHP20k-BC-lipase film after depolymerization has stopped (left) and fluorescent microscope image of the same film (right) demonstrating that lipase is still embedded in the film even after degradation stops. The enzyme was labelled, and the film imaged, using procedures as previously reported.^[10]^ b) Hydrolysis of a small molecule ester by BC-lipase embedded in a PCL-RHP20k film that has stopped degrading, proving that the enzyme has not denatured within the film.

**Figure S6.**
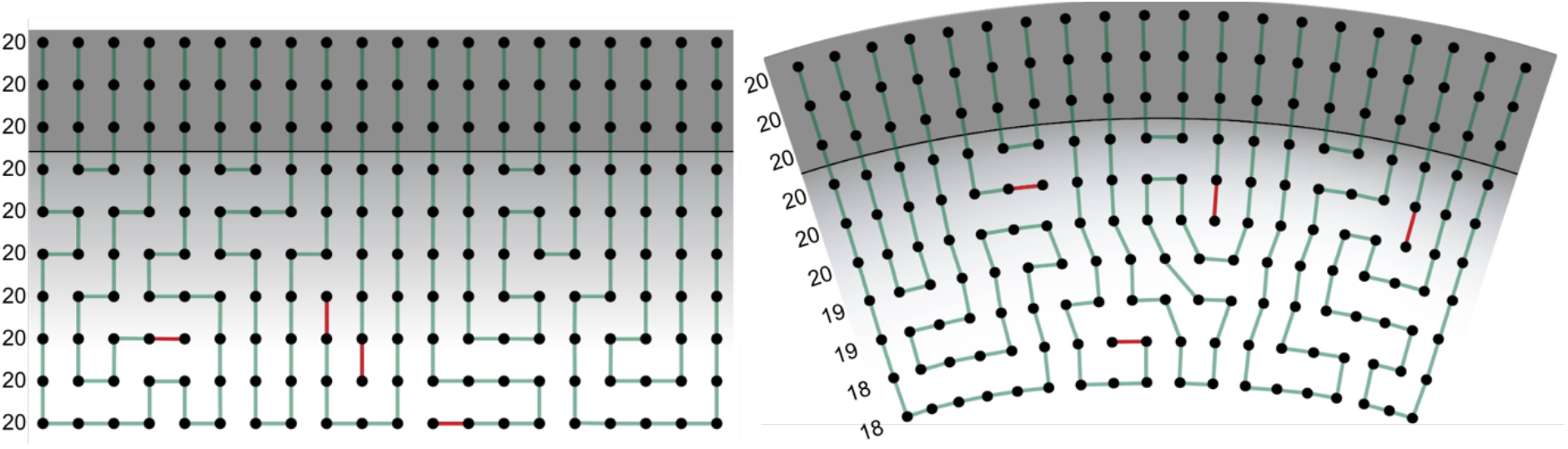
Lattice model of planar (left) and curved (right) lamellae; an adjacent crystalline (top) and amorphous (bottom) domain demonstrates how curvature creates reduced available volume at increasing distances below the crystal-amorphous interface, while a planar morphology possesses constant available volume throughout. The numbers to the left indicate total number of lattice sites available for that layer. Chain end segments are indicated in red, while all other polymer segments are in green. For curved lamellae, the same number of polymer chain segments emerging from the crystalline domains are able to be accommodated in the amorphous domain if more chains terminate near the crystal/amorphous interface. No such constraint exists for planar lamellae.

**Figure S7.**
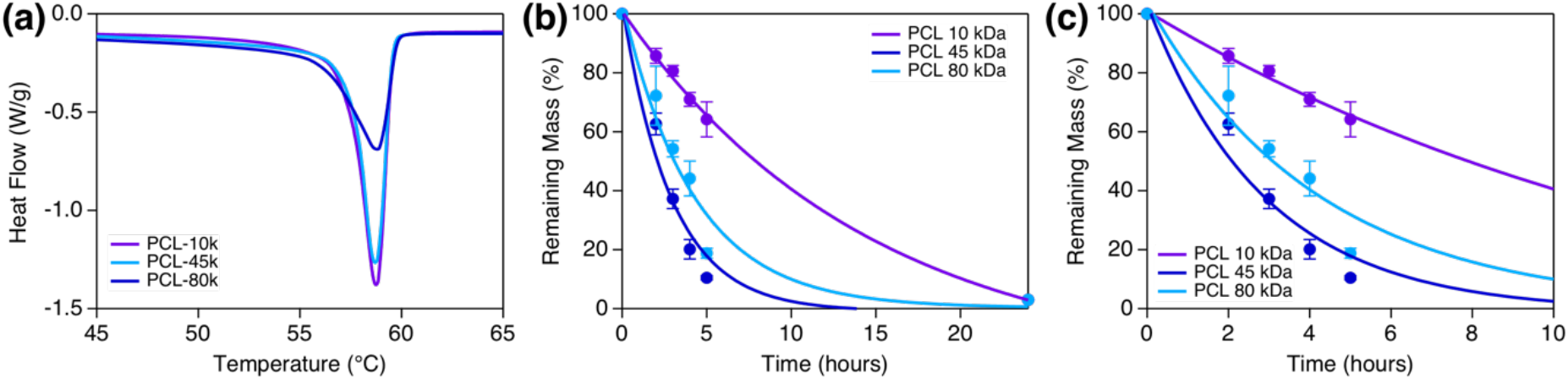
a) DSC of melt-recrystallized PCL films demonstrating similar percent crystallinities and crystalline lamellae thicknesses regardless of PCL MW; b) depolymerization of PCL-RHP-BC-lipase as a function of molecular weight demonstrating that each film degrades to near completion at 37 °C over 24 hours; c) zoomed-in portion of graph S5b highlighting that the lowest MW PCL sample depolymerizes at the slowest rate, suggesting chain end binding is a rate-limiting factor.

**Table S1.** T2 decay values from NMR experiment described in Figure 3e, calculated by fitting a double exponential; error represents the uncertainty of the fit from each experiment.

| Sample | Rigid-amorphous<br>T2 decay [ms] | Mobile-amorphous<br>T2 decay [ms] |
| --- | --- | --- |
| Pure PCL, as cast | 0.18 ± 0.02 | 5.4 ± 0.5 |
| PCL-RHP<sub>70k</sub>-BC-lipase<br>50% degraded | 0.14 ± 0.03 | 3.2 ± 0.3 |
| PCL-RHP<sub>20k</sub>-BC-lipase<br>50% degraded | 0.78 ± 0.17 | 8.1 ± 6.9 |

